# Addition of a single bacterial isolate to conventional larval rearing water can impact wing size and longevity in adult male *Aedes aegypti*

**DOI:** 10.64898/2026.01.27.701390

**Authors:** Camden M. Dezse, Noha K. El-Dougdoug, Sarah M. Short

**Affiliations:** Department of Entomology, The Ohio State University, Columbus, OH, USA; Botany and Microbiology Department, Faculty of Science, Benha University, Benha, Egypt

**Keywords:** *Aedes aegypti*, Cedecea, microbiota, bacteria, life history, longevity, mass rearing

## Abstract

Male mosquitoes are mass-reared around the globe for use in mosquito control programs like Sterile Insect Technique and Incompatible Insect Technique. During larval development, mosquitoes co-exist with complex microbial communities that serve as food and also form the internal microbiota of the organism. The microbiota can impact multiple larval and adult life history traits including development rate and male body size. In the present study, we investigated how male wing length and adult male longevity is impacted by the addition of a single bacterial isolate to otherwise conventional larval rearing water. Of three isolates tested, we found that larval exposure to one, *Cedecea sp*., resulted in adult males with significantly reduced wing length and longevity. Our findings suggest that minor modifications of the microbial community during larval development can have life-long effects on mosquitoes which, like many organisms, acquire their microbiota from the environment. Moreover, they suggest that mosquito life history traits can be influenced by small shifts in the microbial rearing community, which could impact the efficacy and efficiency of mosquito mass rearing for vector control.

## Background

Sterile insect technique (SIT) and Incompatible Insect Technique (IIT) are widely used methods of biological control to manage populations of insects (1–6). In both SIT and IIT, released males mate with wild females and resulting offspring are nonviable, causing populations of targeted insects to decrease (7–10). Historically, this technique has seen great success in the control of the screw-worm fly and other fruit fly pests (11), and multiple versions of SIT/IIT have resulted in successful local control of *Ae. aegypti* (8,12–15). Success of this technique relies in part on efficient and cost-effective mass-rearing of male insects that are able to reliably mate with wild females. Life history traits that may be relevant to the success of SIT/IIT include larval development/pupation rates, synchronization of development times, adult body size, and longevity (16).

The bacterial microbiota, i.e. the bacteria found within and in association with *Ae. aegypti* mosquitoes, has been shown to influence the rate and overall success of development, wing length, and longevity in both males and females (17–22) as well as reproduction and vector competence in females (21–28). Therefore, the microbiota may have the potential to impact the efficiency of mass rearing and the quality of mosquitoes produced by mass rearing facilities (29). However, most of the studies investigating impacts of the microbiota on mosquito traits have focused on female mosquitoes (because of their role in pathogen transmission via blood feeding). It is imperative to better understand the effects of the microbiota on male mosquitoes, because male mosquitoes are used extensively in vector control techniques like SIT and IIT. Moreover, most previous studies have been conducted under conditions where the conventional microbiota was removed, and extraneous microbes were continuously excluded (i.e. monoxenic or gnotobiotic settings). Controlled studies of this nature are extremely valuable for understanding the mechanistic role of the microbiota in mosquito life history traits but may be less relevant in an applied setting where gnotobiotic rearing would be impractical. Additionally, Roman et al., 2024 (30) showed that addition of *Asaia* bacteria to *Ae. aegypti* larval rearing water induced significant changes in pupation rate when the conventional microbiota was intact but not when it was removed, suggesting that applying microbial treatments in the presence of the conventional microbiota has the potential to reveal effects on mosquito life history traits that would otherwise remain hidden.

In the current study, we amended the rearing water of *Ae. aegypti* larvae with one of three bacterial isolates previously cultured from *Ae. aegypti* mosquitoes—*Asaia sp., Cedecea sp., and Kocuria sp*. This was done under non-sterile rearing conditions that allowed for the formation of a conventional microbiota. We then measured wing length—as a proxy for body size—and longevity in adult males that arose from each treatment versus a control group with no microbiota manipulation. We chose these bacterial strains because *Asaia* bacteria have been shown to alter pupation success and development time in *Ae. aegypti* under gnotobiotic and non-gnotobiotic conditions (27,30) as well as male wing length under gnotobiotic conditions (27). *Cedecea* bacteria can form a biofilm in the larval digestive tract and have the capacity to competitively exclude other microbes from colonizing *Ae. aegypti* (31,32). *Kocuria* has not been directly implicated in alteration of mosquito life history traits to our knowledge, but it is a Gram-positive bacteria previously found to be associated with multiple mosquito species (33–35).

## Methods

### Mosquito rearing

We reared *Ae. aegypti* (Liverpool strain) using standard rearing conditions and rabbit blood (Hemostat, USA), as described in MacLeod et al. (2021) (36). The mosquitoes were maintained at 27°C and a relative humidity of 80% on a 14h light: 10h dark cycle.

### Experimental set-up and bacterial treatment of mosquito larvae

The effects of three different genera of bacteria were investigated: *Asaia sp., Cedecea sp*., and *Kocuria sp*. Strains of these genera were previously isolated from *Ae. aegypti* digestive tracts (37,38) and have been maintained in 15% glycerol stocks since isolation. We Sanger sequenced the 16S rRNA gene from each strain using primers fd1 and rp1 (39). Sequences were trimmed and aligned using Unipro UGENE (v 52.1) and submitted to Nucleotide BLAST using default parameters and the nr/nt Nucleotide collection database. This revealed that the best match for *Kocuria sp*. is *Kocuria rhizophila* (partial sequence length = 1382bp, query cover = 100%, percent identity = 100%, E value = 0.0), for *Asaia sp*. is *Asaia krungthepensis* (partial sequence length = 1351bp, query cover = 100%, percent identity = 99.85%, E value = 0.0) and for *Cedecea sp*. is *Cedecea neteri* (partial sequence length = 1403bp, query cover = 100%, percent identity = 100%, E value = 0.0), Supplementary File S1. For each of four independent experiments, four groups were created: a control group consisting of larvae with no bacterial amendment, and three groups of larvae where we amended the larval water with live bacteria from one of the aforementioned genera (Figure 1). Each larval rearing tray was established with 200 first-instar larvae, one liter of reverse osmosis (RO) water, and two pieces of autoclaved orange cat food (each 200-250 mg) (9Lives Indoor Complete dry cat food). The treatment groups then each received an additional 10^8^ Colony Forming Units (CFUs) of one bacterial isolate suspended in 1X PBS, and the control group received a volume of 1X PBS equal to the mean volume of suspended bacteria added to the other treatments. Trays were handled on the open benchtop and the microbiota was allowed to form conventionally, with the exception that bacterial amendments were added to the three experimental trays.

**Figure 1.**
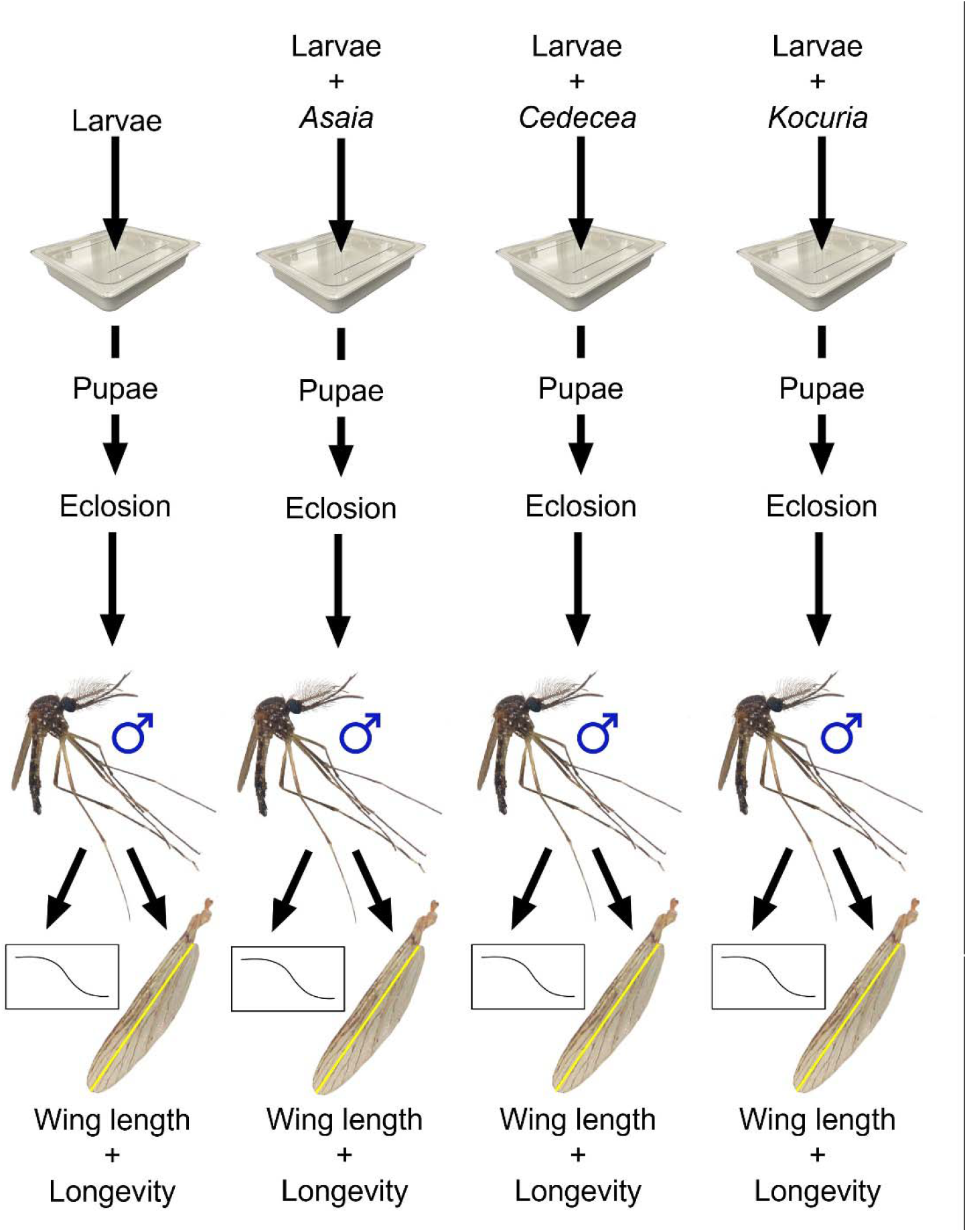
Experimental Design. Four groups were created: a control group consisting of larvae with a conventional, unaltered microbiota, and three groups of larvae with bacteria added to the larval rearing water in addition to the conventional microbiota. Each larval rearing tray received larvae, sterile water, and dry cat food. The treatment groups each received 10^8^ CFUs of a single bacterial isolate suspended in 1X PBS, and the control group received 1X PBS. The larvae from each group developed into pupae and were placed into plastic cages with mesh lids. Three days post peak eclosion, 25 male mosquitoes from each group were placed into small soup cups with mesh lids to measure longevity, and the right wings of all remaining adult male mosquitoes were measured. The full experiment was replicated four times, and total sample sizes were (88-139) for longevity and (35-96) for wing length measurement for each treatment.

Pupae were transferred to cups and placed in large plastic cages with mesh lids. Adults were allowed to feed *ad libitum* from a 10% sucrose solution in a bottle located in each container until three days post peak eclosion, at which point males were separated from females based on standard morphological features (40) and transferred to smaller containers for longevity observation.

### Wing length measurement

Wing length is a measurement used to estimate body size of mosquitoes (40). Using a binocular microscope (Leica S9i) and subsequent analysis in ImageJ v1.53, the right wings of males from each treatment were measured to the nearest 0.01mm. Wings were measured from the distal end of the alula to the tip of the R3 vein (41).

### Longevity measurement

Three days post peak eclosion, all mosquitoes from each container were cold anesthetized and 25 male mosquitoes from each group were placed into pint-size cardboard cups with mesh lids and allowed to feed *ad libitum* on a 10% sucrose solution. Each day, dead individuals were removed until no live individuals remained in the cups.

### Statistical analysis

Statistical analyses were conducted using R statistical software and R Studio (42,43). Because overall wing size varied substantially across replicates (Figure S1), we opted to group mean-center the wing length data by subtracting the average wing length for each replicate from all data points within that replicate.. We then used a two-way

ANOVA with treatment and replicate as the predictor variables followed by pairwise contrasts using Tukey’s methodTo analyze survival data, we first fitted survival curves using the Kaplan-Meier method. In order to assess the effect of treatment on survival, we fit a Cox proportional-hazards model with treatment as a predictor variable. We initially included replicate in the model as well, but the assumption of proportional hazards was not met for the replicate variable, therefore we opted to exclude it from our final model. All raw data can be found in Supplementary File S1. R code and output for wing length and survival data can be found in Supplementary File S2.

## Results

### Effect of bacterial treatment on wing length

The effect of bacterial treatment on the wing lengths of adult male *Ae. Aegypti* was assessed. Treatment had a significant effect on male wing length (Figure 2; F_3,265_ = 6.07, p = 0.00052), and pairwise comparisons via Tukey’s HSD revealed that males from the *Cedecea* treatment had significantly shorter wings than males from the negative control treatment (Figure 2; p = 0.001) while those from the *Kocuria* treatment (Figure 2; p = 0.985) and the *Asaia* (Figure 2; p = 0.596) treatment did not.

**Figure 2.**
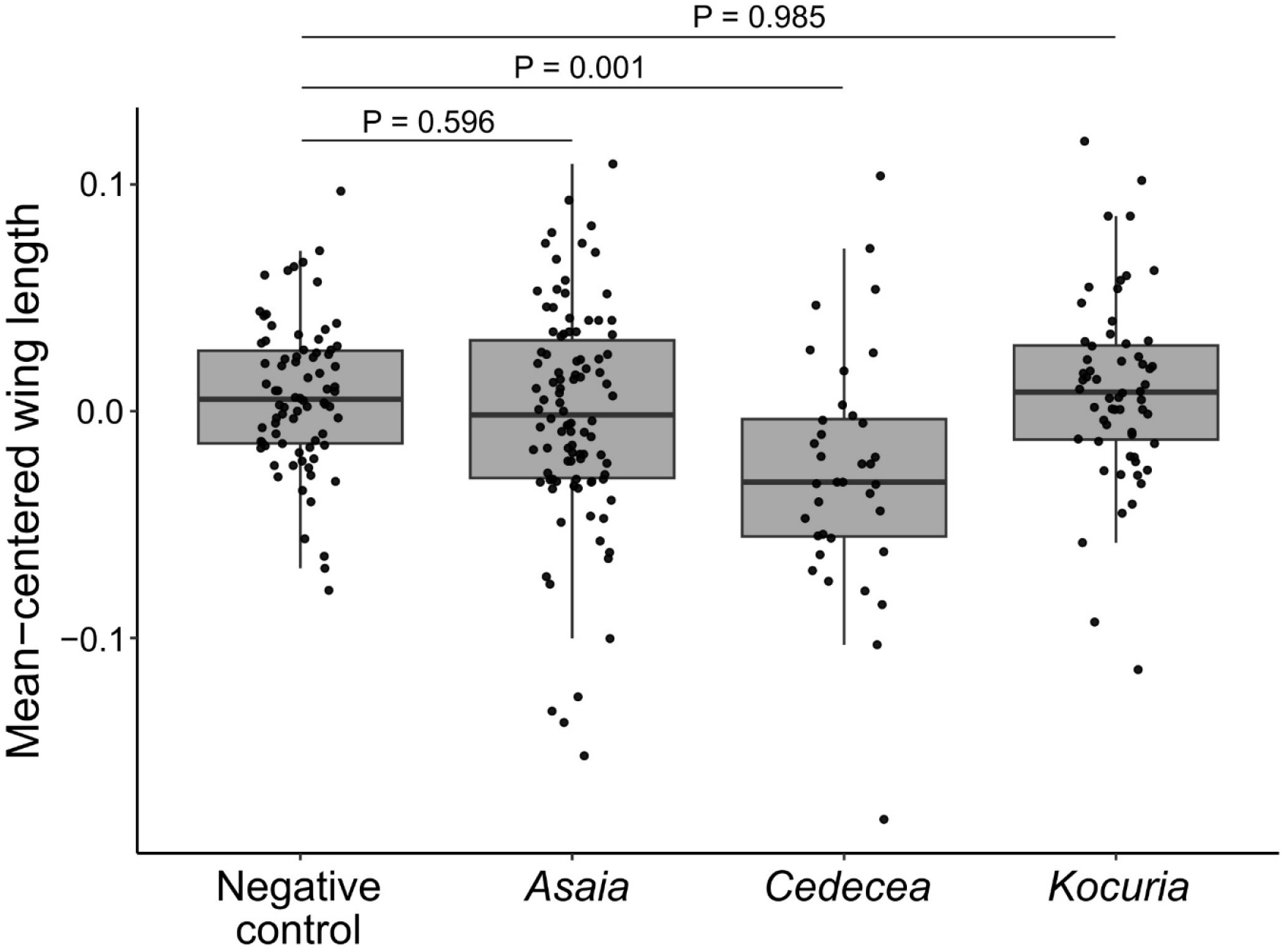
Wing lengths are significantly shorter for adult males from the *Cedecea* treatment compared to the negative control. Right wings of adult males from each treatment were measured from the alular notch to the tip of the R3 wing vein. The experiment was repeated four times. Due to variation in overall wing size between replicates, data were mean-centered within replicate prior to analysis. Points on each box plot represent individual group mean-centered male samples. Treatment significantly impacted wing length (F_3,265_ = 6.07, p = 0.00052). P values indicate significant differences between treatments and the negative control as revealed by Tukey’s HSD. Total sample sizes were as follows: Negative control, n = 78; *Asaia*, n = 98; *Cedecea*, n = 36; *Kocuria*, n = 60.

### Effect of bacterial treatment on longevity

Bacterial treatment significantly affected survival (Figure 3; p = 0.022). Pairwise contrasts indicated that *Cedecea* significantly decreased longevity when compared to the negative control (Hazard ratio_*Cedecea* vs. Neg. ctrl._ = 1.58, p = 0.007), while *Asaia* and *Kocuria* did not (Hazard ratio_*Asaia* vs. Neg. ctrl._ = 1.18, p = 0.25; Hazard ratio_*Kocuria* vs. Neg. ctrl._ = 0.96, p = 0.80).

**Figure 3.**
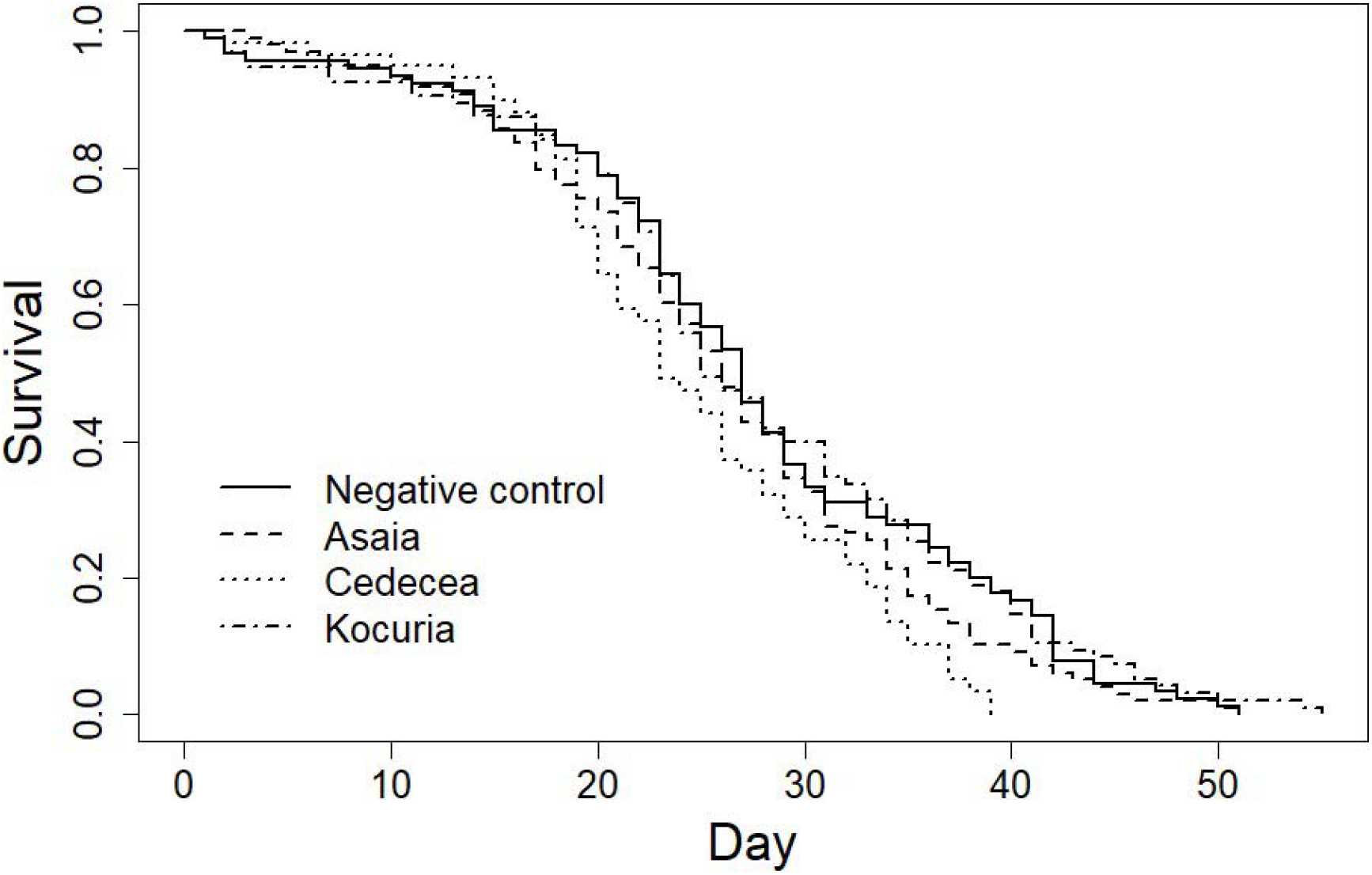
Longevity is significantly reduced for adult males from the *Cedecea* treatment. Three days post peak eclosion, males from each treatment were collected and placed into cups to measure longevity. Resulting data was pooled, fitted using the Kaplan-Meier method, and a Cox proportional hazards model was applied. Bacterial treatment was found to significantly affect survival (LR test = 9.6, p = 0.02), and pairwise comparisons indicated that *Cedecea* significantly decreased longevity compared to the negative control (Hazard ratio_*Cedecea* vs. Neg. ctrl._ = 1.58, p = 0.007). In contrast, neither *Asaia* nor *Kocuria* treatment significantly altered longevity (Hazard ratio_*Asaia* vs. Neg. ctrl._ = 1.18, p = 0.25; Hazard ratio_*Kocuria* vs. Neg. ctrl._ = 0.96, p = 0.80). The experiment was repeated 3-4 times per treatment and total sample sizes were as follows: Negative control, n = 90; *Asaia*, n = 98; *Cedecea*, n = 59; *Kocuria*, n = 95.

## Discussion

We found that amendment of larval water immediately after hatching with bacteria from the genus *Cedecea* had a significant effect on body size and longevity of male *Ae. aegypti*. When compared to a control group receiving no additional bacterial treatment, adult male *Ae. aegypti* from the *Cedecea* treatment group had smaller bodies (estimated via wing length) and decreased longevity. This adds further support to the growing body of evidence that microbes in aquatic habitats impact the life history of mosquito larvae and subsequent adults. Moreover, it suggests that bacterial amendments to conventional larval rearing water can shape male mosquito life history traits even when the microbiota is otherwise allowed to form naturally.

The bacterial microbiota of *Ae. aegypti*, especially during larval development, has been shown to play a key role in shaping life history traits of adult mosquitoes. For example, the larval microbiota has been repeatedly shown to influence adult female (18,20,21,28) and male (18,20) wing length. Adult longevity has also been shown to be impacted by the larval microbiota, however, evidence of this is much more plentiful for females (18,19,21,22,27,28) than for males (19,22). While it is common for mosquito microbiota research to focus on females due to their importance in pathogen transmission, we chose to investigate male mosquitoes because of their use in vector control strategies such as SIT and IIT, both of which utilize the mass rearing and subsequent release of male insects (1–6). Another way in which our study deviated from the majority of previous studies (but see (30)) was in our larval habitat set-up. Most prior studies investigating the impacts of different bacteria on mosquito life history traits have utilized a gnotobiotic rearing system in which the native microbiota was removed and larval water conditioned with a single bacterial isolate and maintained aseptically to ensure monoxenic conditions (17,20,21,27,28). In the current study, we intentionally did not remove the native microbiota from the surface of the eggs and handled the larval trays non-aseptically to determine if addition of bacterial amendments can impact life history traits under non-gnotobiotic conditions.

Our findings suggest that addition of a single bacterial isolate to an otherwise conventional larval rearing environment has the capacity to alter adult male *Ae. aegypti* life history traits. However, only one isolate, *Cedecea sp*., elicited a response, therefore it remains unclear how generalizable this effect may be and further studies on a larger panel of bacterial and/or fungal isolates are warranted. We note that the reduction in wing length as a result of *Cededea* exposure was robustly observed in two replicate experiments; in a third replicate, which was underpowered for the *Cedecea* treatment, we also observed this same relationship between *Cedecea* and the negative control. However, in one replicate, we observed similar wing lengths between the negative control and the *Cedecea-*treated group. Perhaps notably, average wing length was highest in this replicate. This suggests that the effect of *Cedecea* treatment could potentially be condition-dependent, i.e. the effect of *Cedecea* on male life history traits may be most pronounced when males are otherwise stressed.

Another open question is whether the effects of Cedecea would persist without the native microbiota or whether they are dependent on its presence. Roman et al., (2025) (30) found that effects of *Asaia spp*. bacteria on *Ae. aegypti* pupation arose from an interaction with the native microbiota. The bacterial taxa present in the natural larval habitat have been shown to have complex interactions amongst themselves and with developing larvae (e.g. (44)), which further highlights the value of observing microbiota-host interactions in ecologically and practically relevant contexts, rather than solely in gnotobiotic or axenic systems.

Our finding that larval *Cedecea* exposure impacts adult male life history traits is informed by previous research showing that *Cedecea* species form biofilms in the larval digestive tracts of *Ae. aegypti* larvae and competitively exclude other bacterial taxa from colonizing (31,32). This could have significant implications for mosquito life history, especially if *Cedecea* is pathogenic or is suppressing the colonization of microbes that are beneficial to development. Microbes serve as a key source of nutrition for mosquitoes (45) and provide necessary micronutrients, such as B vitamins (46). *Cedecea-*mediated dysbiosis could potentially disrupt beneficial functions of the microbiota leading to smaller males with shortened longevity. Metabolic changes induced in larvae due to interactions with microbes have been seen to affect adult fitness later in development (21), making the aforementioned explanation plausible. However, further study into *Cedecea*-mosquito-microbiota interactions in the context of male life history traits is needed to test this hypothesis.

A key caveat to our study is that we did not evaluate the persistence of *Cedecea* after addition to the larval water during the first instar. It is possible the bacteria failed to persist in the aquatic environment and/or never colonized the larvae themselves. In this case, the effects we detected could have been a result of an early, fleeting larval exposure to the bacteria or an effect of the bacteria on the microbial community which then had subsequent effects on larval development and/or adult male traits. Notably, previous studies have shown that when *Cedecea* is present in the larval water, it is also found in larvae and adults, suggesting that bacteria from this genus have some degree of persistence across developmental stages (36). Nonetheless, further research is warranted to determine how *Cedecea* bacterial amendment of the larval water alters the microbiota of the aquatic habitat and the larvae overall and the extent to which effects on mosquito traits are directly and/or indirectly related to *Cedecea* exposure.

Neither *Asaia* nor *Kocuria* impacted the adult male traits we measured in this study. This is particularly surprising for *Asaia*, given that bacteria from the genus *Asaia* have been shown to alter the pupation success and rate of development in *Ae. aegypti* under gnotobiotic and conventional conditions (27,30), as well as male wing length under gnotobiotic conditions (27). Larvae in our study were not reared gnotobiotically, which may explain why we did not observe an effect on male wing length. Moreover, Roman et al. (2024) (30) showed that *Asaia krungthepensis* (the *Asaia* species most closely related to the strain used in the current study) added to conventional larval rearing water are found in larvae at three days post-hatching but are largely absent by six days post-hatching. Perhaps, therefore, *Asaia* do not persist long enough in a conventional larval rearing environment to elicit an effect on wing length or adult male longevity.

## Conclusion

This study demonstrated that the addition of a single bacterial isolate, *Cedecea sp*., to the larval habitat of *Ae. aegypti* resulted in adult male mosquitoes with reduced wing size and longevity compared to males arising from larval habitats with no bacterial amendments. This suggests that minor changes to the microbial community during larval development can impact mosquitoes well into adulthood. Unlike the majority of research into the mosquito microbiota, which utilizes axenic or gnotobiotic systems, our findings emphasize the profound effects that microbial treatments can have even in the presence of a conventionally formed microbiota. Furthermore, by exclusively focusing on male mosquitoes, our work highlights the potential importance of the microbiota for mass-rearing protocols used in SIT and IIT programs. Expanding this line of inquiry to include additional microbial taxa could further elucidate how microbes affect mosquito development and how those effects may be harnessed to improve vector control.

## Acknowledgements

We wish to thank Dom Magistrado for feedback on analyses and all current and former members of the Short lab for helpful guidance and feedback.

## Funding statement

This work was supported by the National Institutes of Health National Institute of Allergy and Infectious Diseases (grant number R21AI174093), the Ohio State University Infectious Diseases Institute, and the Ohio State University College of Food, Agricultural, and Environmental Sciences. The funders had no role in study design, data collection and analysis, decision to publish, or preparation of the manuscript.

## Data Accessibility Statement

All raw data generated in this study is provided in Supplementary File S1.

## Artificial Intelligence (AI) declaration

ChatGPT-4 was used between the dates of May 2, 2025 and July 16, 2025 to generate initial drafts of portions of the second paragraph of the introduction and the second and third paragraphs of the discussion.

The resulting output was heavily edited after generation and prior to inclusion in the manuscript.

## Supplementary file legends

**Supplementary File S1:** All raw data used in the manuscript, as well as Sanger sequences used to identify the best match taxonomic ID for bacterial isolates.

**Supplementary File S2:** All R code and outputs from all analyses performed in the manuscript.

**Supplementary Figure S1:** Wing length data parsed by replicate. These are the same data used in Figure 2, but here each replicate is plotted separately. Points on each box plot represent individual male samples and data are not mean-centered. Treatment significantly impacted wing length in replicate 1 (F_3,70_ = 3.91, p = 0.012), replicate 3 (F_3,65_ = 6.31, p = 0.0008), and replicate 4 (F_3,89_ = 7.78, p = 0.0001), but not in replicate 2 (F_3,32_ = 1.33, p = 0.28). Letters indicate significant differences between treatments as revealed by Tukey’s HSD within each replicate (p < 0.05). Total sample sizes were as follows: Negative control, n = 78; *Asaia*, n = 98; *Cedecea*, n = 36; *Kocuria*, n = 60

## Notes

### Competing Interest Statement

The authors have declared no competing interest.

