## Supplementary figures and images for "Addition of a single bacterial isolate to conventional larval rearing water can impact wing size and longevity in adult male *Aedes aegypti*"

### Supplementary Figure S1

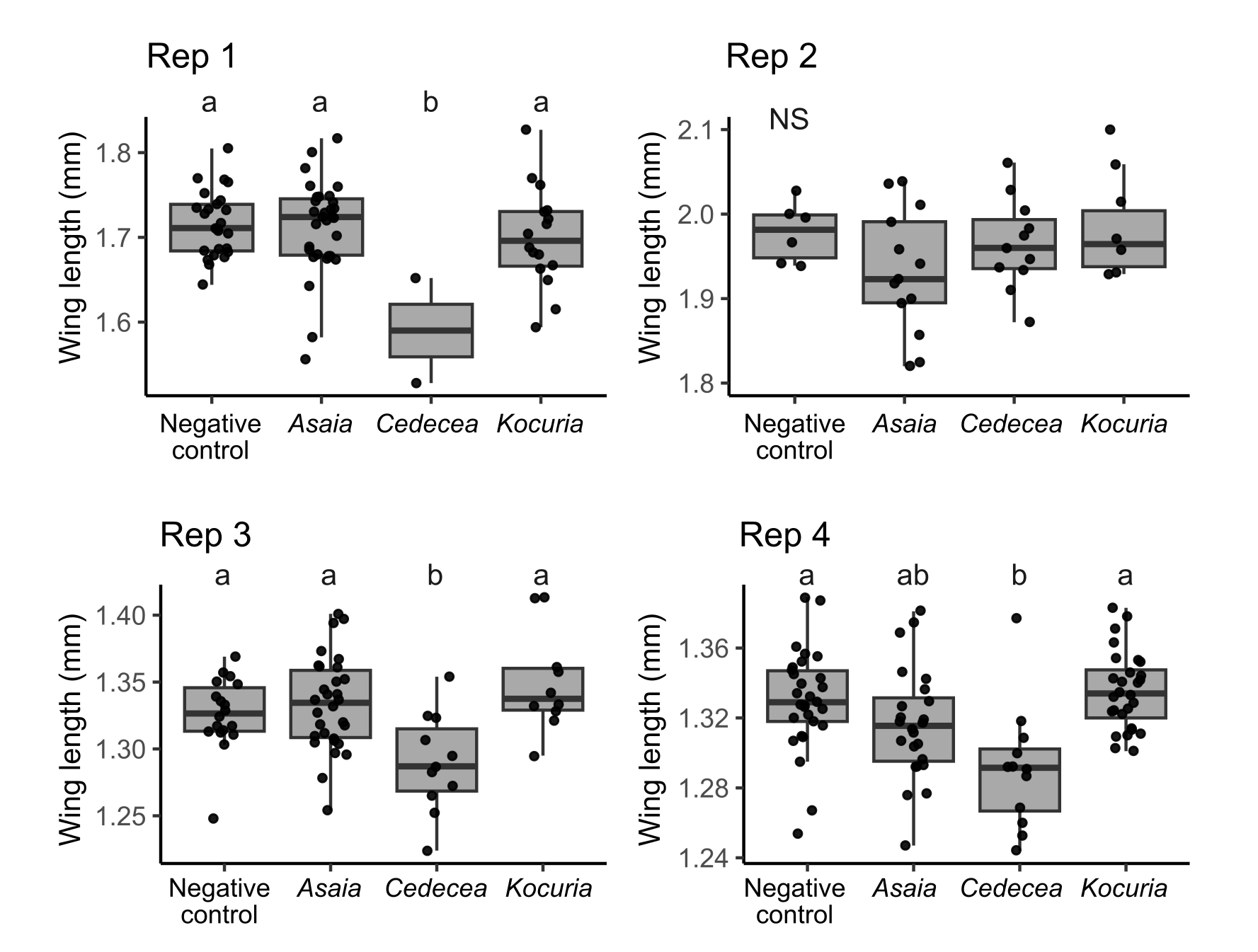
